# Analysis of seven putative Na^+^/H^+^ antiporters of *Arthrospira platensis* NIES-39 using transcription profiling and *In-silico* studies: an indication towards alkaline pH acclimation

**DOI:** 10.1101/344416

**Authors:** Monika M. Jangir, B. Vani, Shibasish Chowdhury

**Affiliations:** Department of Biological Sciences, Birla Institute of Technology and Science, Pilani, Rajasthan, India (333031).

**Keywords:** *Arthrospira platensis* NIES-39, Na^+^/H^+^ antiporters, pH, Real-time PCR, CPA family

## Abstract

Na^+^/H^+^ antiporters mediated pH regulation is one of the known mechanism(s), which advocates a possible role of the antiporters in the alkaline pH tolerance of *Arthrospira platensis* NIES-39 too. Seven putative Na^+^/H^+^ antiporters have been reported in *A. platensis*. We have characterized these seven antiporters, where A1, Q2, L2 and L6 belong to the CPA1 family whereas C5, D5 and O6 belong to CPA2 family through various *In-silico* analysis. Conserved domain analysis of these seven putative antiporters indicate the presence of nine different kinds of domains. Out of these nine domains, six domains function as monovalent cation-proton antiporters and two as the universal stress protein (Usp) category. Transcription profile of these seven antiporters was also generated at three different pH and time frames which showed a significant difference in the mRNA levels at pH 7, 9 and at 11 along with a temporal pattern of the expression profile. The *Insilico* and the real time PCR analysis, put together, suggest the active participation of these seven putative Na^+^/H^+^ antiporters in alkaline pH homeostasis of this cyanobacterial strain where CPA1 subfamily antiporters play a major role.

## Introduction

Ion and pH homeostasis are fundamental regulators of cellular processes. It is reported that among prokaryotic and eukaryotic organisms, pH is regulated by antiporters specially Na^+^/H^+^ antiporters and Mrp-like antiporters [4]. Antiporters maintain a stable pH inside the cell under alkaline conditions [27] and help in the cell volume regulation [24].

All the Na^+^/H^+^ antiporters ranging from prokaryotes to human belong to CPA super family [25]. CPA family is further subdivided in to CPA1 and CPA2 subfamily. The structure and function of Na^+^/H^+^ antiporters was first studied in *E.coli* by Padan et al. [15, 11]. Padan et al. in 1989 studied the role of nhaA gene (Na^+^/H^+^ antiporter) in pH regulation and salt stress by chromosome deletion. Later crystallographic structures of the antiporters along with the functionally important residues became available for other prokaryotes, belonging to CPA1 and CPA2 family.

Further, gene knockout studies of Na^+^/H^+^ antiporters in *Synechocystis* sp. PCC 6803 strongly supported their active participation in pH regulation along with salinity stress [40]. Deletion of plant genes which encodes the vacuolar and cytoplasmic membrane Na^+^/H^+^ antiporters decreases the plant’s salt tolerance [2, 30] whereas over expression is used to produce salt-resistant plants [1].

Many reports thus, suggest the role of Na^+^/H^+^ antiporters in pH regulation. *A. platensis* NIES-39 was isolated from a soda lake, which can withstand the alkaline pH (up to pH 11). Its genome is composed of a single, circular double stranded chromosome of 6.8 Mb, without any plasmids [13]. The mechanism behind pH homeostasis has not been elucidated so far in this organism; however, seven Na^+^/H^+^ antiporters are reported but still putative. Through *In-silico* analysis these seven putative antiporters are compared with the CPA1 and CPA2 family proteins along with their functionally conserved domains. The differential and temporal transcript expression profile of the Na^+^/H^+^ antiporters under different pH regimes was also studied, to appraise their possible role during pH stress.

## Material and Methods

### Sequence analysis and domain search

*A. platensis* NIES-39 antiporter’s protein sequences were retrieved from NCBI database [10]. Pairwise and Multiple sequence alignments for Na^+^/H^+^ antiporters were conducted using ClustalW servers [37]. The domains and motifs of these antiporters were analyzed using the conserved domain (CD) search tools, NCBI [22]. The conservation patterns of the amino acid positions, in the CPA1 and CPA2 family, were analysed using sequence logos [9]. Protein sequences of the *A. platensis* antiporters were retrieved from NCBI and abbreviated as NIES39_A01730 (A1) (WP_014273993.1), NIES39_C00590 (C5) (WP_043468050.1), NIES39_D05350 (D5) (WP_006618914.1), NIES39_L02650 (L2) (WP_006616525.1), NIES39_L06050 (L6) (WP_014276668.1), NIES39_O06850 (O6) (WP_006618272.1), NIES39_Q02770 (Q2) (WP_014277492.1).

### Maintaining Arthrospira platensis NIES-39

*A. platensis* was procured from National Institute for Environmental Studies, Japan, and maintained aseptically in SOT medium (pH 9-9.5) at 30° C temperatures with 4 klux light intensity and a photoperiod of 16h light/8h dark with continues shaking.

### Sample collection at different pH

*A. platensis* was grown at different pH i.e. pH 7, 9 and 11. pH 9 being the optimum pH for the growth whereas 7 and 11 pH were the points used to study pH dependent response of the antiporters. The cells were collected at 0^th^, 1^st^, 4^th^ and 6^th^ hour. Buffered medium was used to maintain a stable pH and the fluctuations were timely monitored.

### RNA Isolation, c-DNA synthesis and real time PCR

The pH treated cells were collected in RNA protect Bacteria Reagent (QIAGEN). Total RNA was extracted using RNeasy mini kit (QIAGEN) and quantified using GE Healthcare Nanodrop. c-DNA was synthesized using Quantitect Reverse Transcription Kit (QIAGEN). iQ SYBR Green Supermix (BIO-RAD) to a final concentration of 1x was used to study the expression profile by semi-quantitative real time PCR. iQ5 multicolour real time PCR detection system from BIO-RAD was used for the PCR. 16s RNA was used as internal control for normalizing the calculations. Primers were designed using Primer3 [38] and the stability parameters were analyzed using IDT Oligoanalyzer 3.1. Primers were synthesized from SIGMA-ALDRICH. The PCR conditions used for the real time PCR starts with an initial denaturation of 3 minutes at 95° C followed by 45 cycles of 95° C-10 seconds, 56° C 30 seconds, 72° C 50 seconds each after which the fluorescence of the product was recorded at 72° C at the end of every cycle. Final extension was given for 10 minutes at 72° C. The calculation of the fold values was done using ^ΔΔ^Ct method [21].

## Results and Discussion

### In-silico characterization of putative antiporters of *Arthrospira platensis* NIES-39

The amino acid sequence length of the seven antiporters from *A. platensis* varies from 409 to 691 (table 1). Pairwise global sequences alignment shows that on an average 17.54% (Standard deviation of 3.58) sequence identity exists among all seven antiporters with maximum 29% sequence identity between D5 and O6 antiporters. Lowest sequence identity of 13.3% is observed between Q2 and D5. Conserved domain database (CDD) search tool identifies nine (among them six are unique) different domains among seven antiporters of *A. platensis* (table 1). Coverage span analysis of each domain reveals that each antiporter of *A. platensis* contains at least one domain belonging to NhaP, Na^+^/H^+^ exchanger, KefB and Asp-A1a exchanger superfamily which exchanges alkali cations such as Na^+^, Li^+^, K^+^, Rb^+^, Ca^2+^ and NH^4+^ against protons [31] across the plasma membrane. Universal stress protein (USP) like domain along with USP A domain, Na^+^/H^+^ antiporter C and TrkA domain along with TrkA-N are other three domains present among the antiporters of *A. platensis*. It has been shown that the expression of USP is enhanced when the bacterial cell is exposed to stress agents [35]. TrkA is a constituent of K^+^ uptake systems and is peripherally bound to the inner side of the cytoplasmic membrane [32]. The presence of all stress related domains within the seven antiporters indicates the possible role of these antiporters in the alkaline stress tolerance mechanism.

**Table 1:** Seven putative Na^+^/H^+^ antiporters and their domain information.

| Antiporters | Abbreviation<br>used for the<br>antiporters | Protein<br>length | Conserved domains | Domain<br>position |
| --- | --- | --- | --- | --- |
| NIES39_A01730 | A1 | 534 | NhaP<br>Na<sup>+</sup>/H<sup>+</sup> Exchanger<br>USP like<br>UspA domain | 1-405<br>5-388<br>420-534<br>418-534 |
| NIES39_C00590 | C5 | 474 | Kef B<br>Na<sup>+</sup>/H<sup>+</sup> Exchanger | 42-473<br>55-457 |
| NIES39_D05350 | D5 | 689 | KefB<br>Na<sup>+</sup>/H<sup>+</sup> Exchanger<br>Na<sup>+</sup>/H<sup>+</sup> AntiporterC<br>USP Like | 10-390<br>16-382<br>422-553<br>563-680 |
| NIES39_L02650 | L2 | 524 | NhaP<br>Na<sup>+</sup>/H<sup>+</sup> Exchanger | 9-426<br>41-395 |
| NIES39_L06050 | L6 | 642 | NhaP<br>TrkA-N<br>TrkA | 1-417<br>404-497<br>404-597 |
| NIES39_O06850 | O6 | 691 | KefB<br>Na<sup>+</sup>/H<sup>+</sup> Antiporter C | 29-415<br>429-557 |
| NIES39_Q02770 | Q2 | 409 | Na<sup>+</sup>/H<sup>+</sup> Exchanger<br>Asp-Ala Exchanger<br>Superfamily | 11-392<br>1-407 |

To further characterize the antiporters, the sequences of seven antiporters were compared with all four prokaryotic Na^+^/H^+^ antiporter whose crystal structures are available. The sequence comparison reveals that all antiporters of *A. platensis* do not have much sequence similarity with *E.coli* Na^+^/H^+^ antiporter (EcNhaA, PDB ID: 1ZCD) which is one of the well-studied antiporter of cation-proton antiporter subfamily 2 (CPA2). PaNhaP, the Na^+^/H^+^ antiporter NhaP from *Pyrococcus abyssi* (PDB ID: 4CZA) [41], has significant similarity with N-terminal domains of Q2 and A1 antiporters whereas MjNhaP1, NhaP1 antiporter from *Methanocaldococcus jannaschii* (PDB ID: 4CZB) [29] is quite similar to the N-terminal domains of L6 and L2. NapA, the antiporter from *Thermus thermophiles* (PDB ID: 4BWZ) [19] has significant similarity with N-terminal domain of C5, D5, O6 antiporters. The domain comparison of *A. platensis* antiporters is given in table 2. It is noticed that Na^+^/H^+^ exchanger domain of around ~400 residues at the N-terminal end of Q2 and A1 antiporters is homologous to PaNhaP, a CPA1 subfamily of cation-proton antiporter which indicates that A1 and Q2 antiporters of *A. platensis* belong to CPA1 subfamily. L2 and L6 antiporters also possibly belong to CPA1 subfamily as N-terminal NhaP domain of these antiporters are homologous to both PaNhaP and MjNhap1 with expectation (E) value of ≤ 1 × 10^-10^ (table 2). KefB domain containing C5, D5 and O6 antiporters belong to CPA2 subfamily antiporters as these antiporters are homologous to NapA with E-value of less than ≤ 1 × 10^-17^. In addition to KefB domain, D5 contains Na^+^/H^+^ antiporter C domain and USP like domain at its C-terminal side whereas O6 contain Na^+^/H^+^ antiporter C domain towards its C-terminal side which accounts for additional ~200 residues in D5 and O6 antiporters.

**Table 2:** *A. platensis* NIES-39’s Na^+^/H^+^ antiporters vs CPA1 and CPA2 antiporters.

| Antiporters | Reference protein | Query coverage | E-value | Identity |
| --- | --- | --- | --- | --- |
| A1 | PaNhaP (4CZ8) | 64% | 3e-14 | 29% |
| Q2 | PaNhaP (4CZ8) | 85% | 2e-16 | 25% |
| L2 | MjNhaP (4CZB) | 63% | 1e-10 | 25% |
| L6 | MjNhaP (4CZB) | 45% | 4e-21 | 25% |
| C5 | NapA (4BWZ) | 88% | 2e-62 | 36% |
| D5 | NapA (4BWZ) | 58% | 2e-17 | 24% |
| O6 | NapA (4BWZ) | 43% | 1e-18 | 25% |

Table 2: Comparison of seven putative Na^+^/H^+^ antiporters from *Arthrospira platensis* NIES-39 against crystal structures of CPA1 (PDB ID: 4CZ8 and 4CZB) and CPA2 (PDB ID: 4BWZ) subfamily.

The amino acid sequence comparison of the seven antiporters of *A. platensis* with available crystals structures (PDB ID: 4CZA and 4CZB) of Na^+^/H^+^ antiporters reveals that the key residues of both CPA1 and CPA2 subfamilies are conserved among these seven antiporters. The detailed analysis of the Na^+^/H^+^ antiporter from Pyrococcus abyssi (PaNhaP, PDB ID: 4CZA) and *Methanocaldococcus jannaschii* (MjNhap1, PDB ID: 4CZB) has revealed that Asp130, Asp159, Thr129 and Ser155 (residues are indicated according to PaNhaP crystal structure numbering) are involved in direct or water mediated indirect ion binding. The oxygen atoms of side chain or main chain carbonyl oxygen atoms are responsible for this protein-substrate ion interaction. The multiple sequence alignment (MSA) of two crystal structures with A1, Q2, L2 and L6 antiporters of *A. platensis* shows that these four residues (except Ser155 in L6 antiporter) are conserved (or replaced with similar amino acids) in CPA1 subfamily indicating prominent role of these residues in substrate ion binding. Both the crystal structures (PDB ID: 4CZA and 4CZB) of CPA1 subfamily have also revealed that Asp93, Thr129, Asn158, Glu154 and Arg337 are part of polar cavity which facilitates ion transport across membrane. Our sequence analysis shows that Arg337 is completely conserved among A1, Q2, L2 and L6 antiporters whereas most of the other polar residues are either conserved or replaced with other polar side chain. Only in case of L6 antiporters, Asn158 was replaced with non-polar residues. Interestingly, even the residues (Ile151, Phe355 and Gly359) whose structural rearrangement is responsible for blocking of ion transportation [41] at low pH are conserved among all CPA1 family antiporters. From the sequence analysis it is clear that both ion binding cavity as well as polar cavity of ion transportation are highly conserved among A1, Q2, L2 and L6 antiporters indicating that these antiporters belong to CPA1 subfamily.

The sequence comparison of NapA (4BWZ) with EcNhaA and Human Nha2 (available crystal structures of other two CPA2 family members) showed only 22.6% and 18.9% sequence identity respectively which possibly indicates that for the function of this class of protein, overall topology and relative orientation of transmembrane helices are more important than sequence conservation. The pairwise sequence alignment of existing crystal structure of bacterial CPA2 antiporters and C5, D5, O6 antiporters resulted into only 20-32% sequence identity with Lys305 conserved among all the three antiporters whereas functionally important Asp156 residue in D5 and O6 is replaced with polar Thr side chain. In case of O6 antiporter the functionally important Asp157 is replaced with polar Asn. The sequence analysis also shows occurrence of a considerable amount of acidic and basic sidechains (13% to 16%) which is also observed as a characteristic feature of CPA2 family protein.

### Transcription profile of seven putative Na^+^/H^+^ antiporters of A. platensis NIES-39

Many studies have reported the positive correlation of the Na^+^/H^+^ antiporter involvement during alkaline stress tolerance mechanism. A similar role of the antiporters could be related in our cyanobacteria. Transcription profile based upon the Real-Time PCR for these seven putative Na^+^/H^+^ antiporters showed a differential temporal expression pattern with a significant enhancement at pH 11 which was recorded as- A1 showed 40-fold; C5, 11-fold; D5, 7-fold; L2, 10-fold; L6, 49-fold; O6, 11-fold; Q2, 14-fold increase (figure 1a-g). After 1 hour of incubation D5 showed a significant expression while antiporters A1, C5, L6, O6, and Q2 were expressed at the 4^th^ hour of incubation. Antiporters L2 and L6 showed their significant expression on the 6^th^ hour of incubation. Exceptionally, L2 showed 14-fold higher expression at pH 7 as compared with the other antiporters. A differential expression was found for different antiporters at pH 7, 9 and 11. We have also observed that the expression of these antiporters under pH stress is temporally regulated which throw light on the actual importance of the seven antiporters in alkaline stress tolerance mechanism.

**Figure 1.**
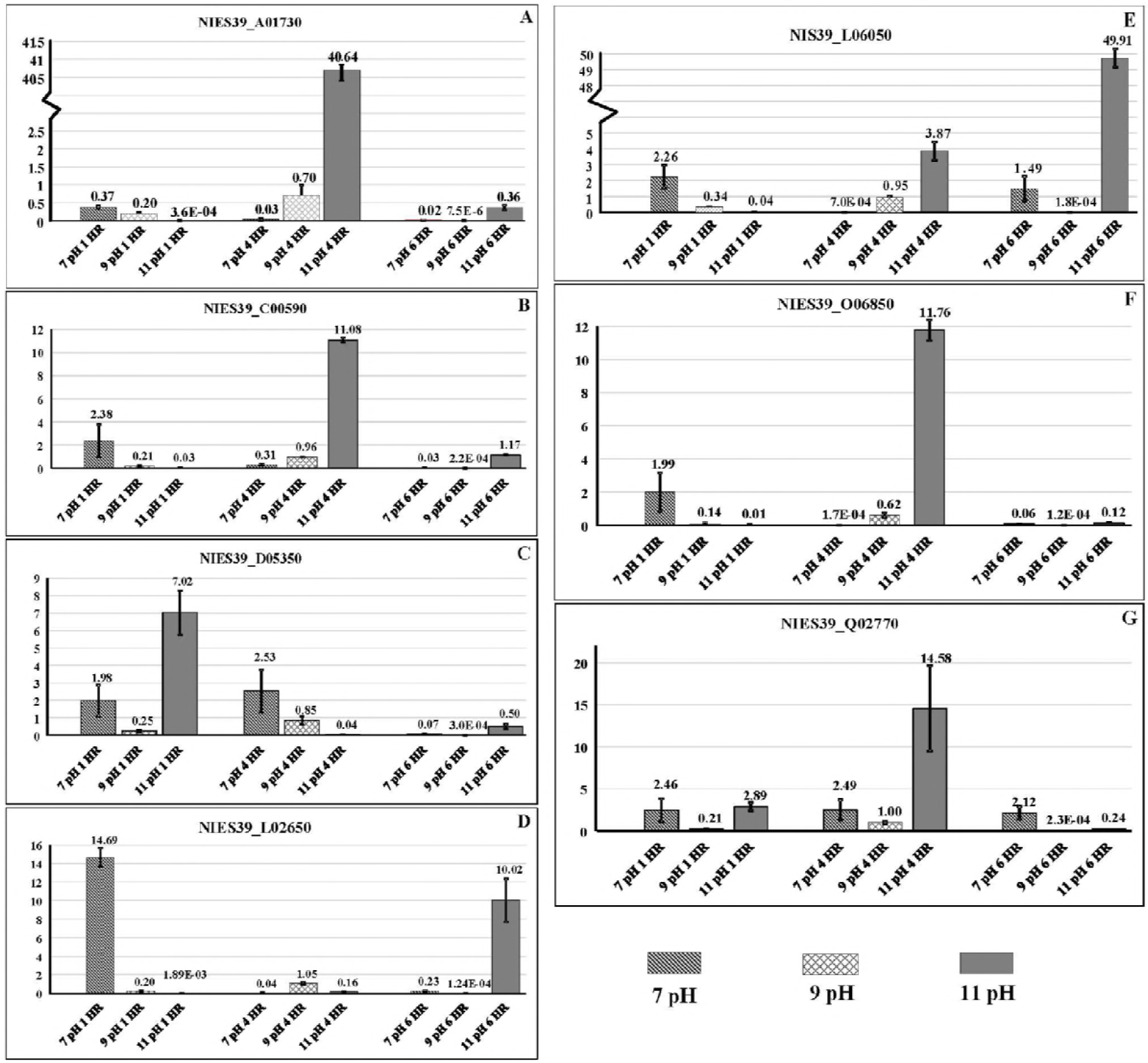
A-E: Expression profile of differential transcripts of A1, C5, D5, L2, L6, O6 and Q2 Na^+^/H^+^ Antiporters from *Arthrospira platensis* NIES-39. Relative gene expression values were normalized using *Arthrospira platensis* NIES-39 16s r-RNA levels as internal loading control. Three biological replicates were taken and the results are mean ± SD of duplicates for each sample.

As seen from our results antiporters, A1 and L6 showed higher level of expression upon subjecting the cyanobacteria to pH 11 as compared to pH 9. Our *In-silico* analysis shows that A1 houses NhaP, Na^+^/H^+^ Exchanger, USP like, Usp A domains and antiporters L6 houses NhaP P, TrkA-N, TrkA domains. Interestingly, only the antiporter, A1 shows the unique UspA domain among the 177 Na^+^/H^+^ antiporters studied from 37 sequenced and annotated cyanobacteria (data not given and to be communicated soon) using various Bio-Informatics tools and databases. Many reports [35, 21] suggest categorically role of UspA in stresses like metabolic, oxidative and temperature. UspA is also found to be associated with pH stress, where its expression is upregulated under alkaline condition in *E. coli* K-12 [42]. Our experiments show that subjecting the cyanobacteria to pH 7 lead to 14-fold higher expressions of the antiporter L2, housing NhaP and Na^+^/H^+^ Exchanger domains (Figure 1d). However, this result could suggest a possible dual role of the antiporter in pH regulation (acidification and alkalization). Also these three antiporters belong to CPA1 superfamily, which suggest an active role played by CPA1 in *A. platensis* as compared to CPA2 superfamily.

## Acknowledgment

The DST-INSPIRE fellowship from DST, India to Monika M Jangir is acknowledged.

## Supporting information

Table 1: The table depicts the 5’ to 3’ base sequences of the forward and reverse primers along with the primer nomenclature for the seven putative Na^+^/H^+^ antiporters from *Arthrospira platensis* NIES-39. 16S rRNA gene was chosen as an internal control.

